# Evaluation of Generative AI Models for Processing Single-Cell RNA-Sequencing Data in Human Pancreatic Tissue

**DOI:** 10.1101/2025.01.15.633192

**Authors:** Sultan Sevgi Turgut Ögme, Nizamettin Aydin, Zeyneb Kurt

## Abstract

Single-cell RNA-seq (scRNAseq) analyses performed at the cellular level aim to understand the cellular landscape of tissue sections, offer insights into rare cell-types, and identify marker genes for annotating distinct cell types. Additionally, scRNAseq analyses are widely applied to cancer research to understand tumor heterogeneity, disease progression, and resistance to therapy. Single-cell data processing is a challenging task due to its high-dimensionality, sparsity, and having imbalanced class distributions. An accurate cell-type identification is highly dependent on preprocessing and quality control steps. To address these issues, generative models have been widely used in recent years. Techniques frequently used include Variational Autoencoders (VAE), Generative Adversarial Networks (GANs), Gaussian-based methods, and, more recently, Flow-based (FB) generative models. We conducted a comparative analysis of fundamental generative models, aiming to serve as a preliminary guidance for developing novel automated scRNAseq data analysis systems. We performed a meta-analysis by integrating four datasets derived from pancreatic tissue sections. To balance class distributions, synthetic cells were generated for underrepresented cell types using VAE, GAN, Gaussian Copula, and FB models. To evaluate the performances of generative models, we performed automated cell-type classification tasks in original and dimensionality-reduced spaces in a comparative manner. We also identified differentially expressed genes for each cell type, and inferred cell-cell interactions based on ligand-receptor pairs across distinct cell-types. Among the generative models, FB consistently outperformed others across all experimental setups in cell-type classification (with an F1-score of 0.8811 precision of 0.8531 and recall of 0.8643). FB produced biologically more relevant synthetic data according to correlation structures (with a correlation discrepancy score of 0.0511) and cell-cell interactions found from synthetic cells were closely resembling those of the original data. These findings highlight the potential and promising use of FB in scRNAseq analyses.

**Author Summary:** Single-cell RNA sequencing (scRNA-seq) analyses focus on identifying distinct cell types and marker genes. Traditional methods face challenges with high dimensionality, sparsity, and sample size imbalances among cell types, limiting automated and unbiased cell-type identification. Generative AI models address these issues by generating synthetic cells for under-represented types, preserving biological and contextual relevance, and employing embedding mechanisms to reduce sparsity and dimensionality. We compared widely used generative models (Variational Autoencoders, GANs, Gaussian Copula, and FB model) using integrated datasets. Synthetic data quality was assessed via cell type classification with Random Forest model in original and reduced feature spaces, correlation of differentially expressed genes, and ligand-receptor interaction inference. The FB model showed the highest potential for creating biologically accurate scRNA-seq profiles. We presented a guideline for automated cell-type identification systems by addressing gaps in single-cell analysis characteristics through the integration of widely used computational biology datasets and generative models (including a novel one, FB model).

## 1. INTRODUCTION

Single-cell RNA-sequencing (scRNAseq) analysis aims to identify different cell types that makeup tissues or tumour microenvironments [1] as well as the marker genes that can distinguish particular cell types from others. Various software platforms and tools have been presented for the implementation and evaluation of these analyses [2,3]. Cell type and marker gene determination usually needs manual operations and it is quite time-consuming. Therefore, in recent years, emphasis has been placed on automating these steps.

Among the analysis tools, Seurat [2] and Single-Cell-Experiment (SCE) [3] R packages stand out and their workflows are similar. Both workflows include; quality control, normalization, feature selection, cell integration, dimension reduction, and visualization steps. After the pre-processing steps, a clustering method (e.g. Kmeans) groups similar cells, and Differential Gene Expression (DGE) is identified for the annotation of cell types. Especially for the annotation purposes, time-demanding work is required in addition to various analyses including manually comparing differentially expressed genes in each cell cluster. So, researchers focused on automation of cell-type identification with low error rates. Comprehensive comparative studies have been conducted for this purpose using machine learning models such as Random Forest, Support Vector Machines, Multi-Layer Perceptrons, and K-Nearest Neighbors, etc. by utilizing datasets from differing species (human, mouse) [4,5], different technologies [6], with sizes, and complexity.

Pre-processing steps importantly affect the downstream processes, i.e. the identification of cell-types and marker genes. Hence in recent years, researchers have been developing frameworks to improve pre-processing steps. Single-cell data consists of large-scale and sparse matrix structures that are computationally challenging to process. Hence the pre-processing step is crucial in terms of affecting the processing time and accuracy of the downstream estimations made. Another problem is the imbalanced class distribution across different cell clusters. Since the validation of findings needs to be carried out in high-cost laboratory environments that contributes to the class imbalance issue, the number of publicly available datasets with annotated and labeled cell types is quite limited [7]. Generative models can address the class imbalance problem by generating synthetic cells, while their embedding mechanisms effectively mitigate the sparsity and high-dimensionality issues [8].

Some studies have utilized the latent space representations of generative models to address the sparsity by reducing the dimensionality. Gronbech et al. [9] proposed a Variational Autoencoder (VAE) based single cell VAE (scVAE) method. This method skips a precise data preprocessing step by using the original count matrix as input and offers a model that can reliably estimate the representations of cells in the latent space. The Gaussian Mixture method, proposed by Choi et al. [10] obtained embedding representations by using the VAE method on both cell and gene basis. Gene-gene relationships and hub genes were identified by creating gene-based embedding representations. On the other hand, generative models have been investigated to address the issues around imbalanced class distributions by generating synthetic cells. Marouf et al. [11] proposed the scGAN model using Generative Adversarial Networks (GANs) for a realistic generation of scRNAseqdata. Their model learns non-linear gene-gene dependencies from complex, multi-cell type samples and uses this information to generate synthetic cells. Yu et al. [12] introduced a novel method called MichiGAN, which combines the strengths of VAE and GAN models. This deep generative model performs sampling using representations that semantically manipulate cells without compromising data quality. Heydari et al. [13] presented the Automated Cell-Type-informed Introspective Variational Autoencoder (ACTIVA) model, which utilizes conditional VAE conditioning on cell-type information during the cell generation process. It consists of three networks; an encoder which is employed as a discriminator distinguishing synthetic cells from real ones, a decoder that works as a generative network creating synthetic data, and a cell-type classification network. ACTIVA demonstrates superior performance compared to solely GAN-based models by generating more realistic synthetic cells. Palma et al. proposed a flow-based generative model named cellFlow [14] utilizing Conditional Flow Matching [15]. This model is based on a likelihood model and negative binomial distribution. The authors highlighted the importance of using raw counts for synthetic cell generation and downstream analysis. They compared the performance of several generative models to cellFlow, finding that cellFlow achieves results comparable to existing methods.

In summary, the structure of single-cell data has urged researchers to adopt generative models into their work. The approaches developed predominantly revolve around VAE, GAN, Gaussian-based models, and more recently, Flow-Based models. Based on this, our study aims to evaluate fundamental generative models as a preliminary work to guide the research community on the development of new single-cell workflows incorporating generative models. Additionally, we seek to highlight the generative capability of relatively more novel and less commonly used Flow-Based models compared to other methods.

## 2. RESULTS

Fig 1 presents our study’s overall framework. Our workflow consists of dataset curation, data preprocessing, dimensionality reduction, synthetic cell generation, and cell-type classification. It also incorporates dataset evaluation, marker gene identification, and cell-cell interaction analysis as key components in downstream analyses.

**Fig 1.**
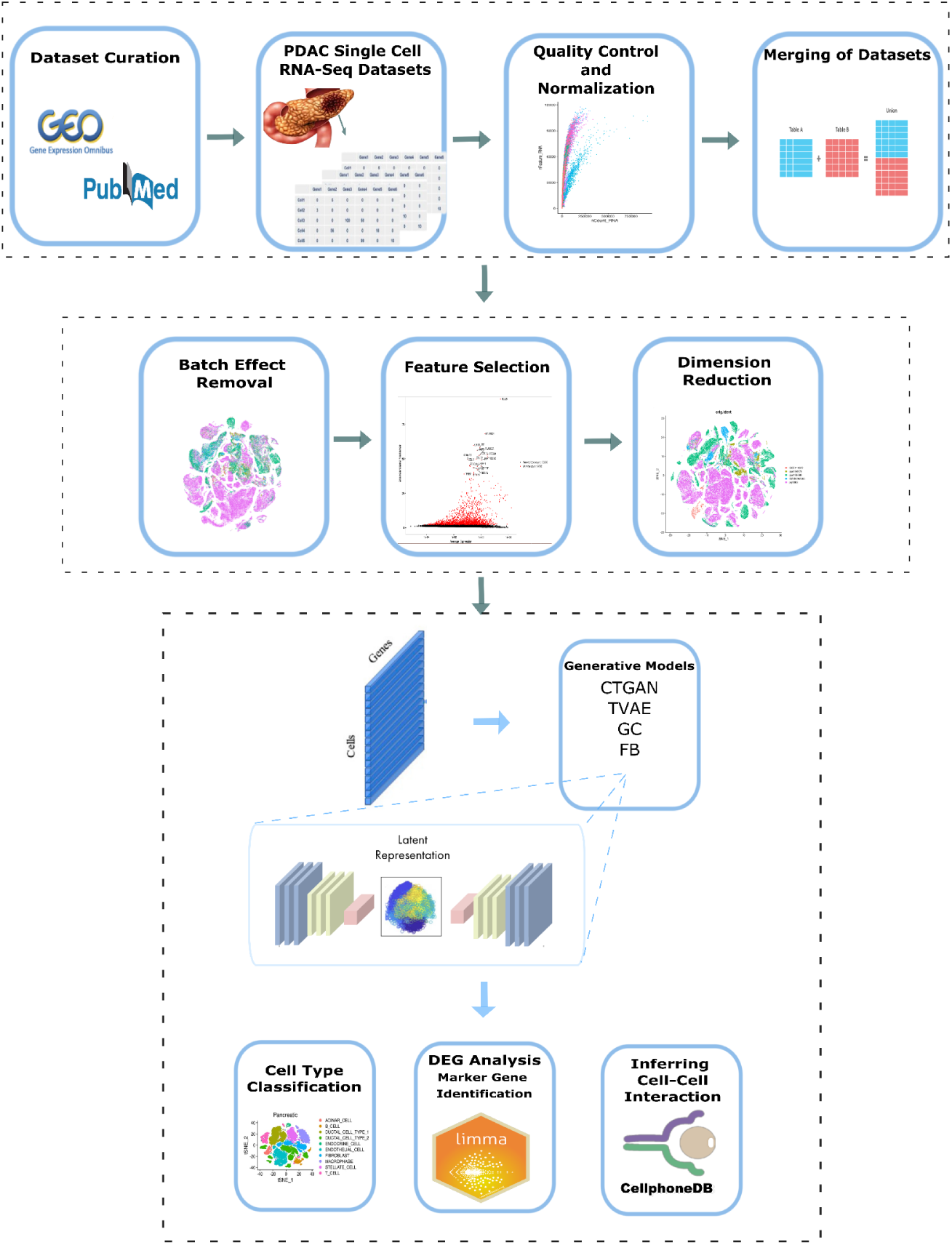
Overall Workflow diagram.

### 2.1. Preparing the Meta Data

Pancreatic tissue datasets are accessible through the R packages *scRNAseq* and the *SingleCellExperiment*. SingleCellExperiment [3] has a similar flow to that of *Seurat*. Firstly, the dataset labels were corrected based on the literature [16]. Classes with less than 10 samples/cells were eliminated and “unclear”, “co-expression”, “not applicable”, and “unclassified” cells were removed from the dataset. Cell types that belong to the same cell category but were incorrectly annotated with different labels were merged. Quality control analysis is applied to each dataset individually as follows: First, genes expressed in less than 100 cells and cells expressing less than 100 genes are filtered out. Cells with very high frequencies (doublets, outliers) are eliminated. Then, the mitochondrial frequency of the cells and the external RNA Controls Consortium (ERCC) were examined and cells with high ERCC values were eliminated. However, it is required to examine these values comparatively, cells with low mitochondrial frequency and high ERCC values may be low quality, while cells with high mitochondrial frequency and low ERCC values may be undamaged active cells. After the quality control step, the datasets are individually normalized using sum factors and logarithmic transformation. The common genes of the datasets are kept to enable meta-analysis for the merged data. Since datasets are curated from different resources, their integration may cause an unintentional bias introduced by the batch effect. Hence, they need to undergo a batch effect correction process. Fig 2 shows the plots of meta-analysis data before and after batch correction performed. Before the batch effect removal, cells obtained from different studies appear separately from one another. But after the batch effect correction, they are more homogeneously distributed on the 2D plane, as represented by the first two principal components of the cells. In the feature selection step, highly variable genes are chosen. The variance of log-expression profiles of each gene is modeled and features are determined based on a suitable mean-variance curve. As a result, 7514 genes were selected from 15839 genes, and the final meta-analysis data has a size of 14172 (cells) x 7514 (genes).

**Fig 2.**
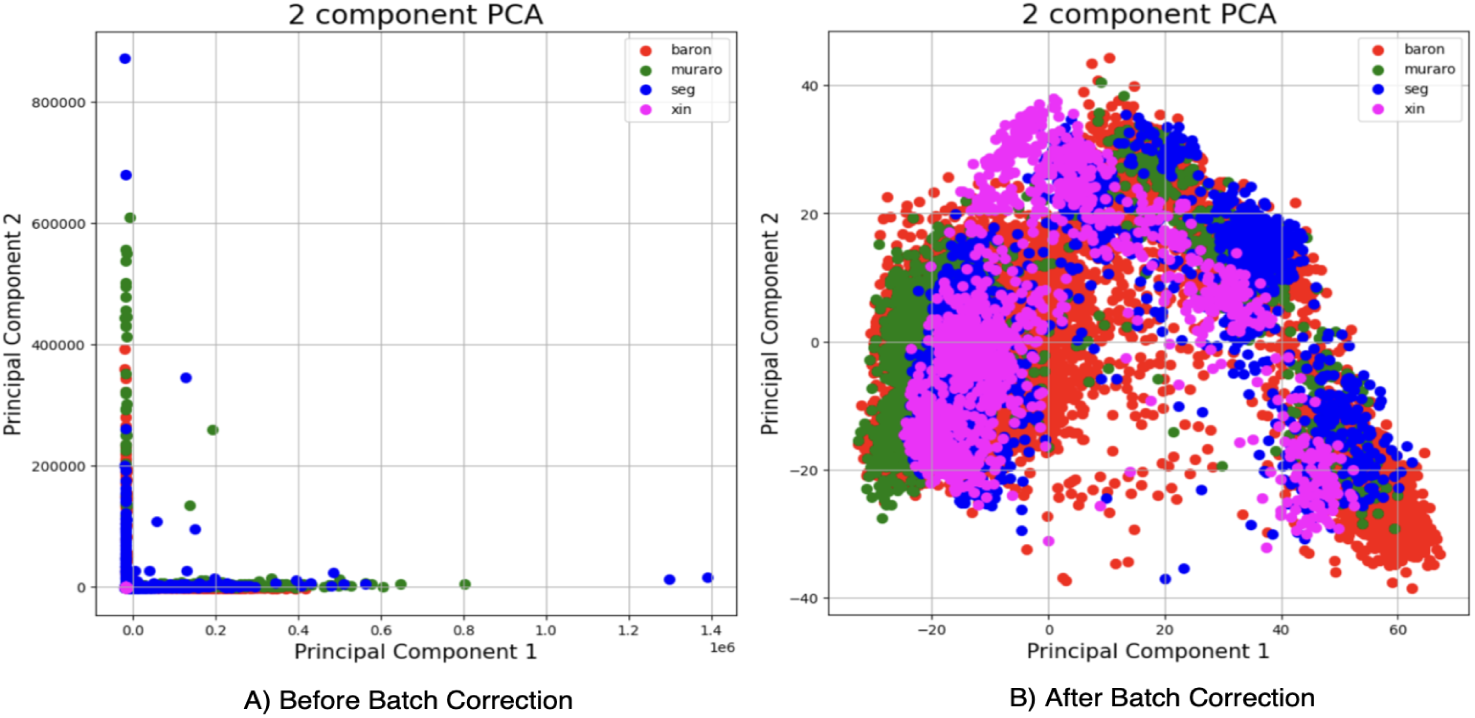
(A) Before and (B) After batch effect removal.

### 2.2. Dimensionality Reduction

We provided inputs to the generative models using three different settings: 7514 data features with no dimensionality reduction, data features reduced to a size of 1500, and data features reduced to a size of 2500. We downsized dimensions of the original data to 1500 and 2500 using PCA. The elbow method was used to determine the optimal dimension size, which returned 2500 (starting point of the curve flattens in x-axis) as the optimum value with a variance of 0.7593(y-axis), and the relevant graph is shown in Fig 3. Additionally, we tested a lower dimension size, 1500, which has an explained variance value of 0.5917 to evaluate generative models under different downsized space conditions.

**Fig 3.**
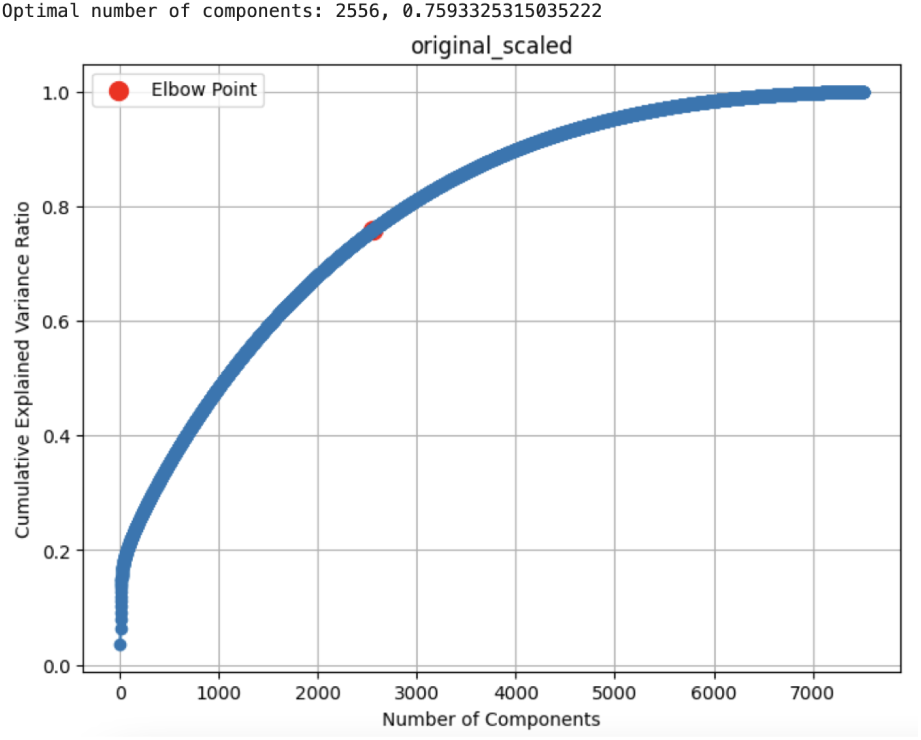
Determining the optimal dimension size for PCA.

### 2.3. Synthetic Cell Generation

To address the class imbalance issue in our training dataset, we applied four generative AI models: TVAE (Tabular Variational AutoEncoder), ctGAN (conditional tabular Generative Adversarial Network), Gaussian Copula (GC), and Flow-Based model, respectively. Each model was used to generate synthetic samples to ensure a more balanced class distribution across all cell types.

We explored the sample size distribution across classes within the training dataset to determine the optimal number of synthetic samples to generate. The third quartile (Q3) value of the sample distribution, which equates to 759, was selected as the target sample size for data generation. For the classes that contain fewer than 759 samples, synthetic cells were generated to reach the number of 759. A minimum sample size of 759 cells in each class was ensured, enhancing the robustness of our model against the class imbalance issue. Fig 4 shows the number of original train samples, synthetic samples and test sets across distinct cell types.

**Fig 4.**
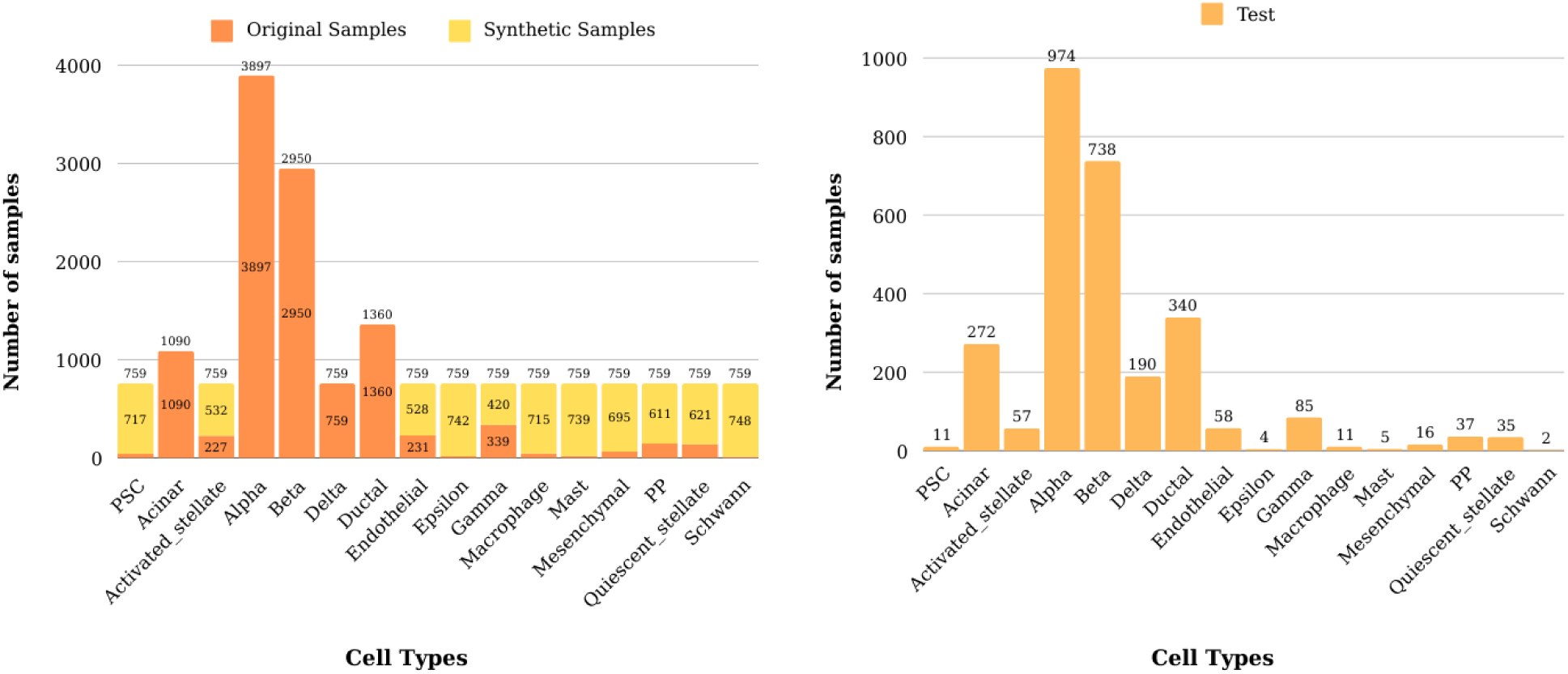
Distribution of cell counts across distinct cell types.

### 2.4. Differentially Expressed Genes (DEG) Identification

Differential Gene Expression Analysis was performed on the test data to identify the top five DEGs associated with each distinct cell type. Among the identified DEGs (adjusted P< 0.05 and log2FC> 1), genes that are mutually exclusive with other cell type categories can be identified as candidate marker genes. [17,18]. For each cell type, violin plots of the top five significant DEGs were generated, and mutually exclusive genes were identified as candidate marker genes for the corresponding cell type. Examples of the generated violin plots are shown in Fig 5. We conducted a literature review to explore previous studies that validated any of the genes listed among the top five DEGs using resources such as DisGeNET [19,20], GeneCards [21,22], and a PubMed search. Some examples of mutually exclusive DEGs that are found in pancreatic tissue, relevant cell-type and associated with tumour include: S100A14 [23], FLT1 [24], PVALP [25,26], RGS5 [27], TPSAB1 [28,29], ITGB2 [30,31], CSF1R [32,33].

**Fig 5.**
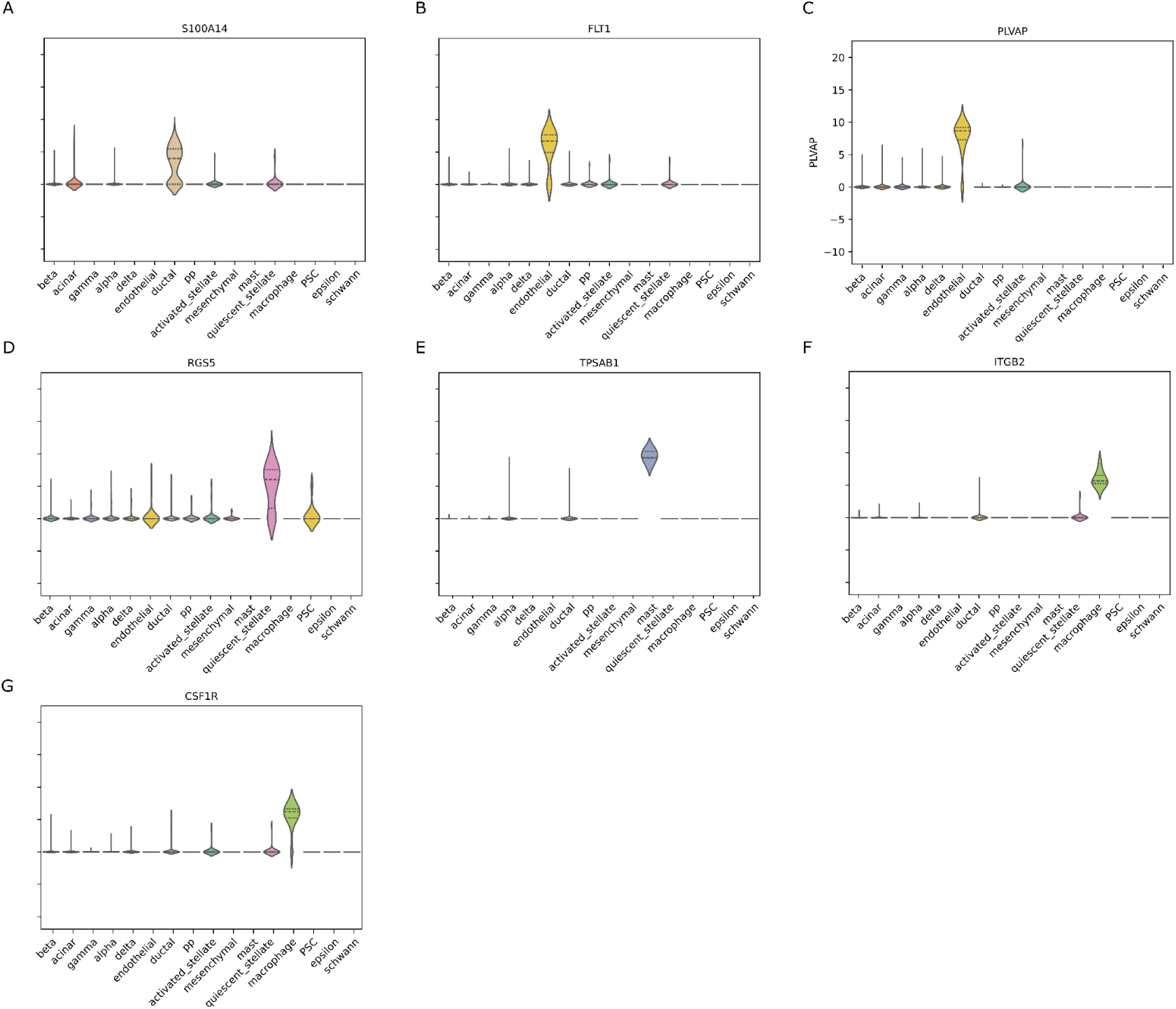
Example of mutually exclusive DEGs found in literature (A) S100A14 (B)FLT1, (C) PLVAP, (D) RGS5, (E) TPSAB1, (F) ITGB2, (G) CSF1R.

### 2.5. Evaluation of Cell Embeddings

The original training samples, test samples, and generated cells (created solely from training samples) were embedded in a 2-dimensional space using PCA. The distribution of samples (original cells plus synthetic cells created using different generative models) on a 2-D scatter plot is shown in Fig 6. Generative models should preserve both biological and contextual patterns present in original data points and retain the relationship motives that exist between original features. So, we evaluated the correlations of the top-5 genes and analyzed the differences between the original and generative sets. To assess this, the Correlation Discrepancy (CD) metric was used, CD calculates the differences in correlation structures. Lower CD values indicate that the generated data has a similar structure to that of the original data. We used the top five DEGs as features while calculating the CD. We avoided using the training data samples, while calculating the CD scores, as the training set’s CD scores would be expected to be close to zero. Instead, we used unseen and unprocessed test set samples for comparison to assess which generative model demonstrates better generalizability. As shown in Table 1, the FB model has the lowest CD value, showing its superiority to the other three generative models in creating real-like synthetic data.

**Fig 6.**
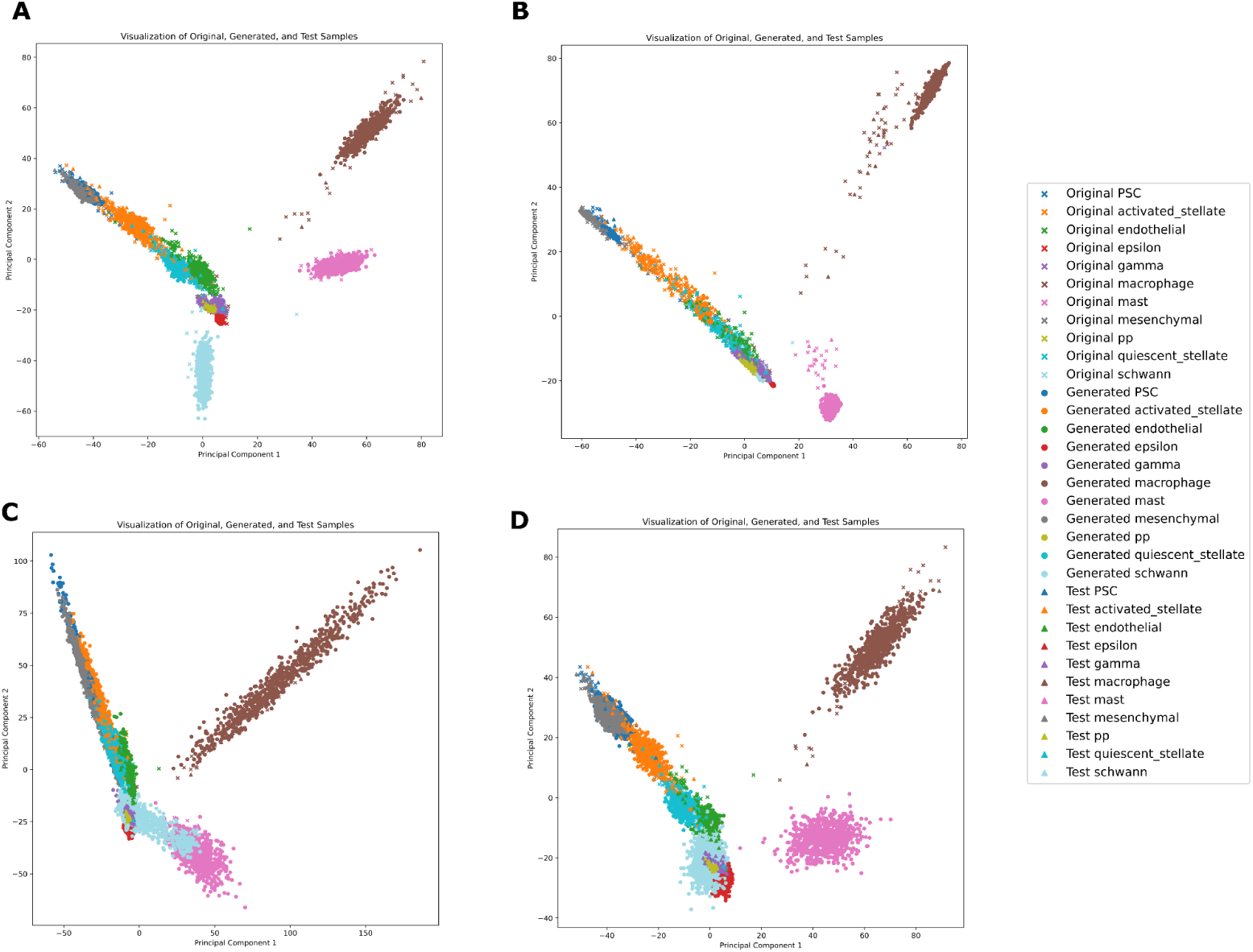
Distribution of original training, generated (synthetic) and test cells in 2-dimensional space where data generation is done by (A) ctGAN, (B) TVAE, (C) GC, (D) FB.

**Table 1.** CD values calculated between the synthetic and test data.

|  | Original Train<br>vs<br>Test | ctGAN<br>vs<br>Test | TVAE<br>vs<br>Test | GC<br>vs<br>Test | FB<br>vs<br>Test |
| --- | --- | --- | --- | --- | --- |
| CD | 0.0149 | 0.0590 | 0.0902 | 0.1602 | **0.0511** |

### 2.6. Cell Type Classification

Cell type classification was performed using the Random Forest (RF) classification model. We trained the RF with solely original data (i.e. real cells) as well as with merging the original (real) data with synthetic cells that are generated by different generative models. Afterward, the RF model was evaluated on a separate test set, which was not used for synthetic data generation to avoid any data leakage, for each experimental setup. Table 2 demonstrates the evaluation results where we have not applied any feature extraction process and used selected highly variable genes among the original features as inputs. The best-performing result is highlighted in bold, while a star next to a result indicates the second-best performance. The results show that while ctGAN demonstrated a slight advantage in the Precision metric, the Flow-Based model achieved the highest overall performance across all metrics.

**Table 2.**
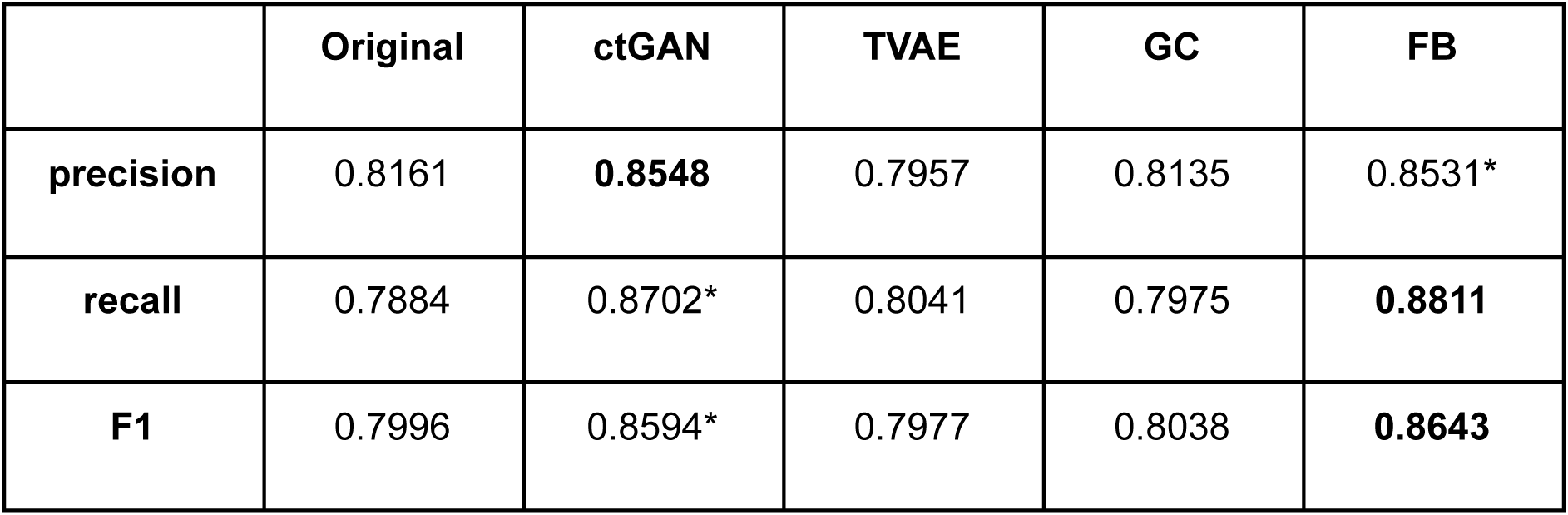
Test results for cell-type classification without performing feature extraction.

|  | Original | ctGAN | TVAE | GC | FB |
| --- | --- | --- | --- | --- | --- |
| **precision** | 0.8161 | **0.8548** | 0.7957 | 0.8135 | 0.8531* |
| **recall** | 0.7884 | 0.8702* | 0.8041 | 0.7975 | **0.8811** |
| **F1** | 0.7996 | 0.8594* | 0.7977 | 0.8038 | **0.8643** |

A secondary approach to comparing generative models is where we generated synthetic cells from the original data, whose dimensionality is reduced to 1500 and 2500, respectively, utilizing PCA. Similarly, the generated cells were combined with the original cells, and cell type classification was performed using the RF model. Fig 7 shows Precision, Recall, and F1-measure results for experimental setups where the first 1500 and 2500 principal components were used as input features. FB performs superior to all other generative models across all evaluation metrics, followed by TVAE.

**Fig 7.**
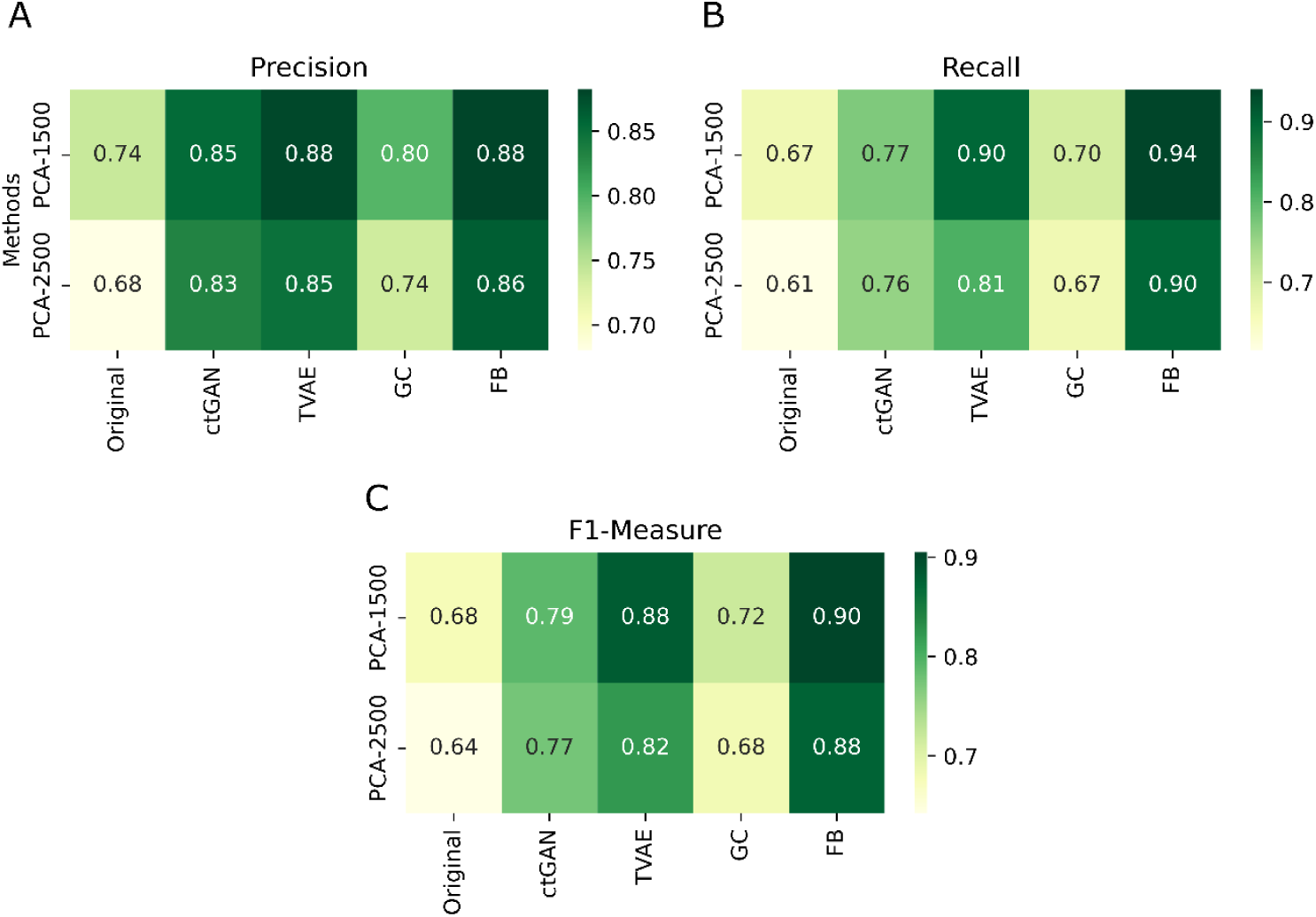
Test performance results for cell-type classification where feature extraction is performed, where the dimension is reduced to 1500 and 2500, respectively (A) precision (B) recall (C) F1-score.

### 2.7. Inferring Cell-Cell Interactions

For inferring cell-cell interactions through identifying potential ligand-receptor communication pairs, CellPhoneDB tool is frequently preferred [34,35]. We investigated the cell-cell interactions in the test, and generative models data. Fig 8 presents heatmaps illustrating the total number of significant cell-cell interactions that exist between different cell types.

**Fig 8.**
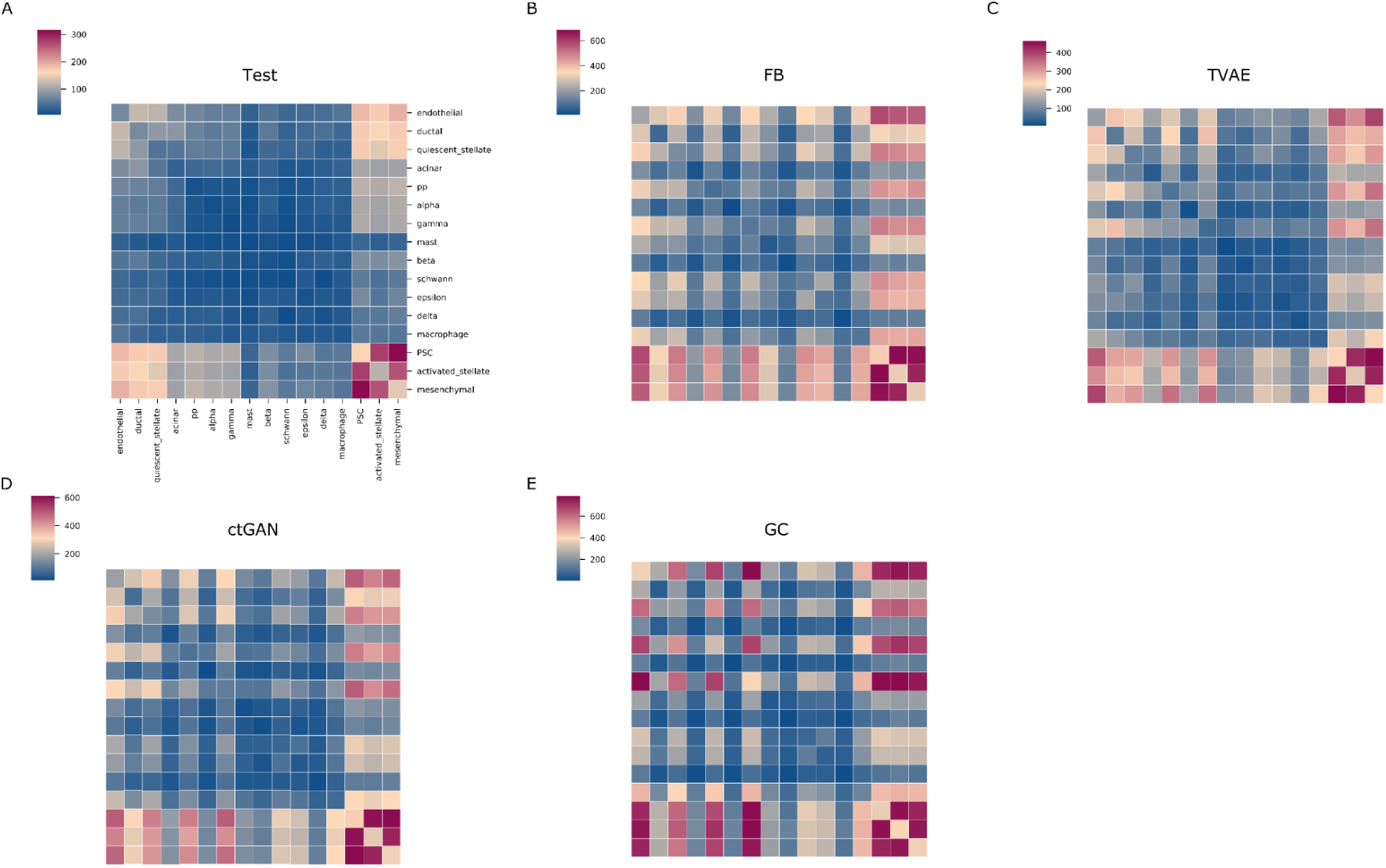
Total number of significant interactions between cell types for (A) Test set, (B) FB, (C) TVAE, (D) ctGAN, (E) GC.

From Fig 8-A (heatmap of cell-cell interaction for the test data),

- Cell type crosstalks between PSC (Pluripotent stem cell)-mesenchymal and PSC-activated stellate show the highest number of interactions.
- Mesenchymal cells interact with cell types PSC, activated stellate, endothelial, quiescent stellate.
- PSC cells interact with: mesenchymal, activated stellate, endothelial, quiescent stellate.
- Activated stellate cells interact with: PSC, mesenchymal, ductal, endothelial.
- Endothelial, ductal and Quiescent stellate cells interact with: PSC, mesenchymal cells.

The cell-cell interaction heatmap for the test set samples is shown in Fig 8-A. Heatmaps for the cells generated by the four generative models (Fig 8 B-E) have higher numbers of interactions but their patterns are similar to the heatmap of the test set, except for the GC model.

Additionally, we discovered ligand-receptor pairs, representing the cell-cell crosstalks, for the test set and generative models. Fig 9 demonstrates ligands, receptors and their interactions for the test set samples. We used the top first most significant DEGs of all cell types for plotting. Due to the large number of interactions between cells, we are unable to show all of the potential ligand-receptor pairs that are identified on a single dotplot. The X-axis refers to cell-cell interaction pairs, the y-axis refers to ligand-receptor pairs. The size of the dots reflects the percentage of the cells in interaction for the corresponding cell types, and the color of the dots, changing from blue to yellow, represents the average of the mean expression value of ligand-receptor pairs in corresponding cells (yellow represents higher expressions). The red outline indicates statistically significant (p < 0.05) ligand-receptor interaction observations. Therefore, it can be assumed that dots with a statistically significant (red outline) representation and a high-valued (yellow) expression are the strongest and most significant interactions.

**Fig 9.**
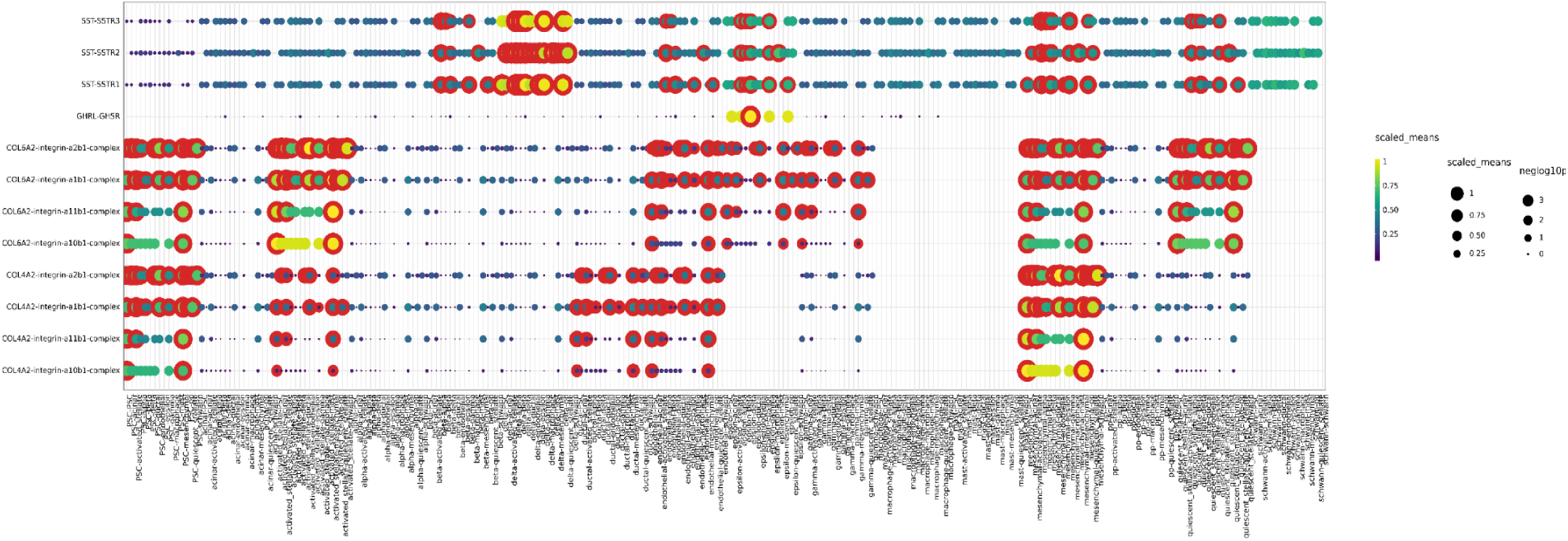
Ligand-receptor pairs representing the cell-cell interactions.

Fig 9 can be summarized as follows;

Ligands correspond to cell type 1, while receptors correspond to cell type 2. First, we listed the cell types associated with the ligands and then identified significantly observed and high-valued (red outline and yellow dots) pairs.

- **COL4A2 (ligand) - integrin-a10b1-complex (receptor), a11b1-complex (receptor), a1b1-complex (receptor), a2b1-complex (receptor)**: Ligands listed here belong to mesenchymal and PSC cell types. Significant and high-valued interactions (red outline and yellow dots) exist between the mesenchymal-PSC pair, mesenchymal-endothelial pair, and mesenchymal-mesenchymal pair.
- **COL6A2 (ligand) - integrin-a10b1-complex (receptor), integrin-a11b1-complex (receptor), integrin-a1b1-complex (receptor), integrin-a2b1-complex (receptor)**: Ligands listed here belong to PSC, activated stellate, mesenchymal, and quiescent stellate cell types. Significant and high-valued interactions (red outline and yellow dots) exist between the activated stellate-PSC pair, activated stellate-endothelial pair, and activated stellate-mesenchymal pair.
- **GHRL (ligand) - GHSR (receptor):** Represent significant and high-valued interactions (red outline and yellow dots) between the epsilon-delta cell type pair.
- **SST (ligand) - SSTR1 (receptor), SSTR2 (receptor), SSTR3 (receptor):** Represent interactions between delta cells and others including activated-stellate and PSC cells.

Dot plots of generative models (S1 Appendix) are similar to those of the test set, with the exception of GC. Generative models have a higher number of interactions overall. However, the significant and high-valued cell-type interaction pairs remain consistent with the test. We observed that the heatmaps shown in the Fig 8 are in line with dot plots representing the strong interactions between cell types PSC, mesenchymal, and activated stellate cells, and so on.

## 3. DISCUSSION

Preprocessing steps in scRNAseq analysis are crucial for identifying cell types, because single cell data is sparse, has large-dimensionality and imbalanced class distributions. Although there are various publicly available scRNAseq datasets in the literature, certain cell types are under-represented and form a minority category, so especially the automated identification of these cell groups is challenging. To address these issues, generative models have been proposed and used in recent years. The VAE model has been widely employed in several studies [9,13]. Additionally, GANs and Gaussian-based models have also been explored [11,36]. Flow-based models have very recently been applied to single-cell analysis studies and indicated a promising potential. We conducted a comparative study by using the most prominent and promising generative models within a well-defined workflow. Our aim was to provide a preliminary pilot study that can serve as a reference guidance for developing new, more robust, and automated cell type identification workflows that benefit from generative models to address class imbalance, sparsity, and high-dimensionality issues.

We conducted a meta-analysis by merging commonly used four scRNAseq datasets [4–6,37] including cell type annotations with a focus on pancreatic tissue, which has been rarely examined. Subsequently, we used four generative models to balance the number of samples across different classes by increasing the sample sizes of minority classes. Synthetic cells generated by each generative model were combined with the training data. The purpose of generative models is to produce synthetic data which is similar to the original one. We demonstrated distributions of training, test, and generated cells along with their cell type annotations in Fig 6. While distinct cell types appear to be separable across all models, the 2-D plot of the FB model (Fig 6-D) shows more distinct separation between different cell types, as expected for the original cells. To validate this observation, the Correlation Discrepancy (CD) metric was used for measuring the discrepancy between train vs test set (as a ground-truth) samples and generated (synthetic) vs test set samples. A smaller CD value indicates higher similarity and the FB model achieved the smallest CD value, 0.0511 (Table 1), implying that it generates samples most similar to the original test data.

The RF, which is a supervised classification model, was trained for cell type classification. Trained models were tested on a previously unseen test dataset. As given in Table 2, the FB model outperformed other methods, followed by the ctGAN model. TVAE and GC models lagged behind. Furthermore, one of the key steps in single-cell analyses is the dimensionality reduction step, where many scRNAseq-analyser tools utilize PCA. Similarly, we reduced the dimensions of the original data from 7514 to 1500 and 2500, respectively, using PCA to evaluate the performance of generative models in the realistic data generation task. As shown in Fig 7, the FB model demonstrates superiority in terms of precision, recall, and F1-measure over other methods. TVAE performed better when the dimensionality reduction is performed, while other methods are less effective with dimensionality reduction when compared to the case of using original, i.e. high-dimensional, input features.

Original single-cell data contains further biological insights, so it is expected that the synthetic data can indicate biologically similar and meaningful outputs. To assess this, cell-cell interactions were inferred and analyzed. In Fig 8, heatmaps of cell-cell interactions were generated using ligand-receptor pairs for test and generative model sets. Among the generative models, the number of cell-cell interactions is similar to those of the test set, except for the GC model. To validate these outputs, ligand-receptor pairs were analyzed and cell-cell interactions inferred between cell types (e.g. Fig 9 and S1 Appendix) were found to be consistent with interactions shown in the heatmaps (Fig 8).

As a result, synthetic cells produced by generative models were analyzed in terms of their cell type classification performances, feature associations, and ability to describe the cell-cell interactions. Overall, the FB model is a promising technique that is suggested to be considered and included in newly developed single-cell analysis tools while addressing computational issues common to scRNAseq processes (i.e. class imbalance, sparsity and high-dimensionality). Consequently, we presented a guideline for a generative model-based automated cell-type identification, which is our contribution to the knowledge and is open to further improvement and developments. In the future, we also aim to explore pancreatic cancer scaRNA-seq datasets to identify interactions between and potential biomarkers of distinct cell types that make up the tumor microenvironment. Pancreatic cancer calls for a better understanding due to its prevalence, poor prognosis and low survival rates [38,39]. We also plan to extend application of our pipeline to scaRNA-seq data from other cancer types or health conditions.

## 4. MATERIALS AND METHODS

### 4.1. Datasets

Relatively limited attention is paid to pancreatic tissue in the literature compared to other tissues. To conduct a comprehensive meta-analysis through integrating various datasets, the human pancreas scRNAseq datasets were curated from sources including GEO Repository [40]. Among the datasets frequently used in computational studies, Baron [4], Muraro [5], Segerstolpe [6], and Xin [37] were selected for our meta-analysis. These datasets, which are appropriate for a preliminary work while assessing generative models, were selected as they possess labels/annotations for cells, have a reasonable size, and are widely used. Table 3 displays the technologies and the cell counts.

**Table 3.** Datasets.

| Dataset | Technology | Cell |
| --- | --- | --- |
| Baron et al year | inDrop | 8569 |
| Muraro | CEL-Seq2 | 2126 |
| Seg | Smart-Seq2 | 2038 |
| Xin | SMARTer | 1600 |
| Total |  | 14333 |

### 4.2. Dimensionality Reduction

We used PCA [41], which is one of the most commonly used dimensionality reduction methods. The basic idea underlying the PCA is to find the first principal component with the largest variance in the data and then search for the second component that is uncorrelated with the first component and explain the next largest variance in the same way, and proceed with the identification of the next components similarly. This process is repeated until the new component becomes almost ineffective in terms of contributing to explaining variance in the original data. It consists of the following steps: standardizing the data, creating a covariance matrix, eigenvalue decomposition, and representing the data in the latent space. PCA is commonly used for both dimension reduction and visualization. We used the PCA function from the Python *scikit-learn* library. The optimal number of PCs is determined using an elbow plot. Components with highest variance values are kept, so the cumulative sum of the explained variance ratios (shown in the y-axis) is calculated for each number of components (x-axis) in the elbow plot. The elbow point (or knee point), where the starting point of the curve flattens and has most of the variance without redundant components, is selected as the optimal PC number. We created the elbow plot with *KneeLocator* function from the Python *kneed* library.

### 4.3. Class Imbalance Issue in scRNAseq Data

In scRNAseqdata, there is usually an imbalance in sample sizes of distinct cell type categories. There are various approaches to eliminate this problem. Synthetic sample generation is an approach to creating new synthetic samples for classes with fewer samples. The main purpose of generative models is to capture the underlying pattern and structure of the training data and create new synthetic data points that are statistically similar to the original data.

Patki et al. [42] created a synthetic data generation python library called Synthetic Data Vault (SDV) for processing tabular data. It is designed to learn complex dependencies of features in datasets and create synthetic data that preserves these dependencies. Generative models are mostly used for creating images; however, the SDV library was particularly developed to facilitate the use of generative models on tabular data. This library includes ctGAN, TVAE, and GC models and it has not been applied to scRNAseq datasets before. To the best of our knowledge, our study is the first to apply this library to a scRNAseq dataset.

Generative models were employed with the training data, while separate test data was reserved for downstream analyses and cell type classification. This approach ensured that there was no data leakage from the test set during the synthetic data generation process.

#### 4.3.1. Conditional Tabular GAN (ctGAN)

ctGAN is a GAN model specifically designed for tabular data. The model learns patterns of the features from the original dataset and generates synthetic data samples that preserve these patterns. GANs [43] consist of two networks that compete with each other: one is a generative network that takes a random noise as input and generates new samples that look like samples from the dataset, and the other one is a discriminator network that tries to distinguish between the real and generated (synthetic) samples. Throughout this comparative training process, the overall model tries to generate more realistic and diverse samples. ctGAN was trained with 50 iterations and other parameters were kept at default values.

#### 4.3.2 Tabular Variational Autoencoder (TVAE)

TVAE is a VAE model designed for tabular data. VAE models learn to encode input data into a lower-dimensional latent space and then generate new examples by randomly decoding data points from this space. VAE models learn a distribution representing the patterns in the dataset and then generate new samples using these patterns learned. TVAE was run with 50 iterations and other parameters were kept at default values.

#### 4.3.3 Gaussian Copula (GC)

GC is a statistical technique used for modeling the dependency structure between variables. Copula models the marginal distributions of individual variables independently and then learns the dependency structure between these independent variables using Gaussian distribution. GC was run with 50 iterations and other parameters were kept at default values.

#### 4.3.4 Flow-Based Model (FB)

FB generative models estimate the exact likelihood of data points, and learn to map data to a latent space and vice versa using invertible transformations. The basic idea of these models [44] is to map a simple distribution, usually a Gaussian distribution, to the target data distribution as close as possible. By applying a series of reversible transformations, these models can capture the gene-gene (i.e. feature) dependencies in the data. *nflows* [45] python library was used and MaskedAffineAutoregressiveTransform [46] was selected as a transformation module. We utilized the method presented in [47] for implementation. For hyperparameter optimisation, combinations of the number of layers {1,2,3,4}, the number of hidden features {128, 256, 512, 1024}, and the learning coefficients {1e-2, 1e-4, 1e-6} were tested. The optimal number of layers was identified as 1, the number of hidden features as 1024, the learning coefficient as 1e-6, and the model was executed through 50 iterations.

### 4.4. Cell Type Identification

We used labeled/annotated data to overcome the class imbalance issue. Therefore, the RF, which is a frequently preferred and employed supervised learning model [25,26], was used. A comparative study was conducted to evaluate the classification performance of the experimental setups where synthetic cells generated by four generative models in both the original feature and dimensionality-reduced spaces.

RF is defined as a forest of decision trees. Decision trees place samples in a hierarchical tree structure from the root node to the leaves. The RandomForestClassifier function was used by using *sklearn* library, and the gain criterion parameter was set to “entropy”. The “n\_estimators” parameter, which specifies the number of trees in the forest, was set to 100. The “max\_depth” parameter specifies the maximum depth of the tree and was set to “none”, which is the default value.

### 4.5. Performance Metrics

We evaluated the cell-type classification performance of cells generated by generative models using precision, recall, and F1-score metrics. Additionally, to assess the correlation between synthetic and original cells, Correlation Discrepancy metric [13] was utilized.

Precision: A metric that measures the ratio of accurately predicted positive samples to all samples predicted as positive. TP: true positives, FP: false positives.

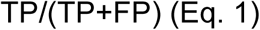

Recall: Measures the ratio of accurately predicted positive samples to all actual positive samples. FN: false negatives

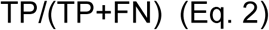

F1-measure: A metric that strikes a balance between precision and sensitivity. F1-measure is calculated using the harmonic mean of precision and recall values.

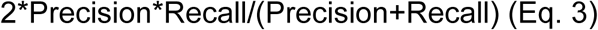

Accuracy: The ratio of correctly predicted samples to the total number of samples.

TN: true negatives

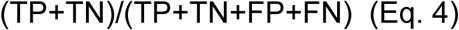

Correlation Discrepancy (CD): measures pairwise correlation matrices of two sets of data by using Pearson correlation, represented by R. Then, it calculates the differences of two correlation matrices and gets their average value. We used this metric to assess the similarity of synthetic (generated) samples to the original ones.

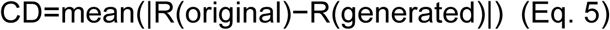

### 4.6. Differentially Expressed Genes Identification

Differential Gene Expression Analysis is performed to detect genes expressed differently across distinct cell types. DEGs demonstrate statistically significant differences in gene expression profiles between cell types. DEGs were detected using Seurat R-package with *findAllMarkers* function [50]. Statistical test parameter was selected as LR (Logistic Regression). The statistical thresholds are adjusted P-value < 0.05 and average log2 Fold Change(log2FC) > 1. We identified the top 5 DEGs for each cell type to evaluate the differences in correlations between features for original and generated cells [13].

### 4.7. Inference of Cell-Cell Interactions and Ligand–Receptor Pairs

Inferring cell-cell communications is essential for understanding several biological processes [28,29]. CellphoneDB is a publicly available tool that identifies ligand-receptor interactions. Cell-cell interactions between cell types are predicted using the Python package CellPhoneDB v5.0.0 [51] with default parameters.

## References

1. Kumar MP, Du J, Lagoudas G, Jiao Y, Sawyer A, Drummond DC, et al. Analysis of Single-Cell RNA-Seq Identifies Cell-Cell Communication Associated with Tumor Characteristics. Cell Rep. 2018;25: 1458–1468.e4. doi:10.1016/j.celrep.2018.10.047

2. Dictionary learning for integrative, multimodal and scalable single-cell analysis | Nature Biotechnology. [cited 27 Nov 2024]. Available: https://www.nature.com/articles/s41587-023-01767-y

3. Amezquita RA, Lun ATL, Becht E, Carey VJ, Carpp LN, Geistlinger L, et al. Orchestrating single-cell analysis with Bioconductor. Nat Methods. 2020;17: 137–145. doi:10.1038/s41592-019-0654-x

4. Baron M, Veres A, Wolock SL, Faust AL, Gaujoux R, Vetere A, et al. A Single-Cell Transcriptomic Map of the Human and Mouse Pancreas Reveals Inter- and Intra-cell Population Structure. Cell Syst. 2016;3: 346–360.e4. doi:10.1016/j.cels.2016.08.011

5. Muraro MJ, Dharmadhikari G, Grün D, Groen N, Dielen T, Jansen E, et al. A Single-Cell Transcriptome Atlas of the Human Pancreas. Cell Syst. 2016;3: 385–394.e3. doi:10.1016/j.cels.2016.09.002

6. Segerstolpe Å, Palasantza A, Eliasson P, Andersson E-M, Andréasson A-C, Sun X, et al. Single-Cell Transcriptome Profiling of Human Pancreatic Islets in Health and Type 2 Diabetes. Cell Metab. 2016;24: 593. doi:10.1016/j.cmet.2016.08.020

7. Wang S, Li H, Zhang K, Wu H, Pang S, Wu W, et al. scSID: A lightweight algorithm for identifying rare cell types by capturing differential expression from single-cell sequencing data. Comput Struct Biotechnol J. 2024;23: 589–600. doi:10.1016/j.csbj.2023.12.043

8. Fajardo VA, Findlay D, Jaiswal C, Yin X, Houmanfar R, Xie H, et al. On oversampling imbalanced data with deep conditional generative models. Expert Syst Appl. 2021;169: 114463. doi:10.1016/j.eswa.2020.114463

9. Grønbech CH, Vording MF, Timshel PN, Sønderby CK, Pers TH, Winther O. scVAE: variational auto-encoders for single-cell gene expression data. Bioinformatics. 2020;36: 4415–4422. doi:10.1093/bioinformatics/btaa293

10. Choi Y, Li R, Quon G. siVAE: interpretable deep generative models for single-cell transcriptomes. Genome Biol. 2023;24: 29. doi:10.1186/s13059-023-02850-y

11. Marouf M, Machart P, Bansal V, Kilian C, Magruder DS, Krebs CF, et al. Realistic in silico generation and augmentation of single-cell RNA-seq data using generative adversarial networks. Nat Commun. 2020;11: 166. doi:10.1038/s41467-019-14018-z

12. Yu H, Welch JD. MichiGAN: sampling from disentangled representations of single-cell data using generative adversarial networks. Genome Biol. 2021;22: 158. doi:10.1186/s13059-021-02373-4

13. ACTIVA: realistic single-cell RNA-seq generation with automatic cell-type identification using introspective variational autoencoders | Bioinformatics | Oxford Academic. [cited 20 Dec 2024]. Available: https://academic.oup.com/bioinformatics/article/38/8/2194/6531957

14. Palma A, Richter T, Zhang H, Dittadi A, Theis FJ. CELLFLOW: A GENERATIVE FLOW-BASED MODEL FOR SINGLE-CELL COUNT DATA. 2024.

15. Tong A, Fatras K, Malkin N, Huguet G, Zhang Y, Rector-Brooks J, et al. Improving and generalizing flow-based generative models with minibatch optimal transport. arXiv; 2024. doi:10.48550/arXiv.2302.00482

16. Tran HTN, Ang KS, Chevrier M, Zhang X, Lee NYS, Goh M, et al. A benchmark of batch-effect correction methods for single-cell RNA sequencing data. Genome Biol. 2020;21: 12. doi:10.1186/s13059-019-1850-9

17. Identification of new marker genes from plant single-cell RNA-seq data using interpretable machine learning methods - Yan - 2022 - New Phytologist - Wiley Online Library. [cited 21 Dec 2024]. Available: https://nph.onlinelibrary.wiley.com/doi/full/10.1111/nph.18053

18. Xu Q, Chen S, Hu Y, Huang W. Single-cell RNA transcriptome reveals the intra-tumoral heterogeneity and regulators underlying tumor progression in metastatic pancreatic ductal adenocarcinoma. Cell Death Discov. 2021;7: 1–16. doi:10.1038/s41420-021-00663-1

19. DISGENET: Genomics Platform for Precision Medicine. [cited 21 Dec 2024]. Available: https://www.disgenet.com

20. Piñero J, Ramírez-Anguita JM, Saüch-Pitarch J, Ronzano F, Centeno E, Sanz F, et al. The DisGeNET knowledge platform for disease genomics: 2019 update. Nucleic Acids Res. 2020;48: D845–D855. doi:10.1093/nar/gkz1021

21. GeneCards - Human Genes | Gene Database | Gene Search. [cited 21 Dec 2024]. Available: https://www.genecards.org/

22. Safran M, Rosen N, Twik M, BarShir R, Stein TI, Dahary D, et al. The GeneCards Suite. In: Abugessaisa I, Kasukawa T, editors. Practical Guide to Life Science Databases. Singapore: Springer Nature; 2021. pp. 27–56. doi:10.1007/978-981-16-5812-9_2

23. Zhu H, Gao W, Li X, Yu L, Luo D, Liu Y, et al. S100A14 promotes progression and gemcitabine resistance in pancreatic cancer. Pancreatol Off J Int Assoc Pancreatol IAP Al. 2021;21: 589–598. doi:10.1016/j.pan.2021.01.011

24. Chung GG, Yoon HH, Zerkowski MP, Ghosh S, Thomas L, Harigopal M, et al. Vascular endothelial growth factor, FLT-1, and FLK-1 analysis in a pancreatic cancer tissue microarray. Cancer. 2006;106: 1677–1684. doi:10.1002/cncr.21783

25. The role of PLVAP in endothelial cells | Cell and Tissue Research. [cited 21 Dec 2024]. Available: https://link.springer.com/article/10.1007/s00441-023-03741-1

26. Khan ST, Ahuja N, Taïb S, Vohra S, Cleaver O, Nunes SS. Single-Cell Meta-Analysis Uncovers the Pancreatic Endothelial Cell Transcriptomic Signature and Reveals a Key Role for NKX2-3 in PLVAP Expression. Arterioscler Thromb Vasc Biol. 2024;44: 2596–2615. doi:10.1161/ATVBAHA.124.321781

27. Zhang S, Fang W, Zhou S, Zhu D, Chen R, Gao X, et al. Single cell transcriptomic analyses implicate an immunosuppressive tumor microenvironment in pancreatic cancer liver metastasis. Nat Commun. 2023;14: 5123. doi:10.1038/s41467-023-40727-7

28. Ye C, Ren S, Sadula A, Guo X, Yuan M, Meng M, et al. The expression characteristics of transmembrane protein genes in pancreatic ductal adenocarcinoma through comprehensive analysis of bulk and single-cell RNA sequence. Front Oncol. 2023;13. doi:10.3389/fonc.2023.1047377

29. Miraki Feriz A, Khosrojerdi A, Erfanian N, Azarkar S, Sajjadi SM, Shojaei MJ, et al. Targeting the dynamic transcriptional landscape of Treg subpopulations in pancreatic ductal adenocarcinoma: Insights from single-cell RNA sequencing analysis with a focus on *CTLA4* and *TIGIT*. Immunobiology. 2024;229: 152822. doi:10.1016/j.imbio.2024.152822

30. Liu X, Rosenthal SB, Meshgin N, Baglieri J, Musallam SG, Diggle K, et al. Primary Alcohol-Activated Human and Mouse Hepatic Stellate Cells Share Similarities in Gene-Expression Profiles. Hepatol Commun. 2020;4: 606–626. doi:10.1002/hep4.1483

31. Benesch MG, Wu R, Menon G, Takabe K. High beta integrin expression is differentially associated with worsened pancreatic ductal adenocarcinoma outcomes. Am J Cancer Res. 2022;12: 5403–5424.

32. Zhu Y, Knolhoff BL, Meyer MA, Nywening TM, West BL, Luo J, et al. CSF1/CSF1R Blockade Reprograms Tumor-Infiltrating Macrophages and Improves Response to T-cell Checkpoint Immunotherapy in Pancreatic Cancer Models. Cancer Res. 2014;74: 5057–5069. doi:10.1158/0008-5472.CAN-13-3723

33. Frontiers | Targeting the CSF1/CSF1R signaling pathway: an innovative strategy for ultrasound combined with macrophage exhaustion in pancreatic cancer therapy. [cited 21 Dec 2024]. Available: https://www.frontiersin.org/journals/immunology/articles/10.3389/fimmu.2024.1481247/full

34. Khaliq AM, Erdogan C, Kurt Z, Turgut SS, Grunvald MW, Rand T, et al. Refining colorectal cancer classification and clinical stratification through a single-cell atlas. Genome Biol. 2022;23: 113. doi:10.1186/s13059-022-02677-z

35. Zhu C, Chen Z, Wang S, Cao J, Cheng Y, Zheng M. Single-cell analyzing of tumor microenvironment and cell adhesion between early and late-stage lung cancer. Mol Immunol. 2024;171: 1–11. doi:10.1016/j.molimm.2024.04.013

36. Sun L, Wang G, Zhang Z. SimCH: simulation of single-cell RNA sequencing data by modeling cellular heterogeneity at gene expression level. Brief Bioinform. 2023;24: bbac590. doi:10.1093/bib/bbac590

37. Xin Y, Kim J, Okamoto H, Ni M, Wei Y, Adler C, et al. RNA Sequencing of Single Human Islet Cells Reveals Type 2 Diabetes Genes. Cell Metab. 2016;24: 608–615. doi:10.1016/j.cmet.2016.08.018

38. Klein AP. Pancreatic cancer epidemiology: understanding the role of lifestyle and inherited risk factors. Nat Rev Gastroenterol Hepatol. 2021;18: 493–502. doi:10.1038/s41575-021-00457-x

39. Blackford AL, Canto MI, Dbouk M, Hruban RH, Katona BW, Chak A, et al. Pancreatic Cancer Surveillance and Survival of High-Risk Individuals. JAMA Oncol. 2024;10: 1087–1096. doi:10.1001/jamaoncol.2024.1930

40. Edgar R, Domrachev M, Lash AE. Gene Expression Omnibus: NCBI gene expression and hybridization array data repository. Nucleic Acids Res. 2002;30: 207–210. doi:10.1093/nar/30.1.207

41. Principal Component Analysis for Special Types of Data | SpringerLink. [cited 20 Dec 2024]. Available: https://link.springer.com/chapter/10.1007/0-387-22440-8_13

42. Patki N, Wedge R, Veeramachaneni K. >The Synthetic Data Vault. 2016 IEEE International Conference on Data Science and Advanced Analytics (DSAA). 2016. pp. 399–410. doi:10.1109/DSAA.2016.49

43. Creswell A, White T, Dumoulin V, Arulkumaran K, Sengupta B, Bharath AA. Generative Adversarial Networks: An Overview. IEEE Signal Process Mag. 2018;35: 53–65. doi:10.1109/MSP.2017.2765202

44. Dinh L, Krueger D, Bengio Y. NICE: Non-linear Independent Components Estimation. arXiv; 2015. doi:10.48550/arXiv.1410.8516

45. Durkan C, Bekasov A, Murray I, Papamakarios G. nflows: normalizing flows in PyTorch. Zenodo; 2020. doi:10.5281/zenodo.4296287

46. Masked Autoregressive Flow for Density Estimation. [cited 21 Dec 2024]. Available: https://www.researchgate.net/publication/317040628_Masked_Autoregressive_Flow_for_Density_Estimation

47. GitHub - uhh-pd-ml/flow-exercise: Exercise on Normalizing Flows, as conducted in the 2nd Terascale School of Machine Learning (2021). [cited 21 Dec 2024]. Available: https://github.com/uhh-pd-ml/flow-exercise

48. Quist J, Taylor L, Staaf J, Grigoriadis A. Random Forest Modelling of High-Dimensional Mixed-Type Data for Breast Cancer Classification. Cancers. 2021;13: 991. doi:10.3390/cancers13050991

49. Xu Y, Li H-D, Lin C-X, Zheng R, Li Y, Xu J, et al. CellBRF: a feature selection method for single-cell clustering using cell balance and random forest. Bioinformatics. 2023;39: i368–i376. doi:10.1093/bioinformatics/btad216

50. FindAllMarkers function - RDocumentation. [cited 21 Dec 2024]. Available: https://www.rdocumentation.org/packages/Seurat/versions/5.0.3/topics/FindAllMarkers

51. Troulé K, Petryszak R, Prete M, Cranley J, Harasty A, Tuong ZK, et al. CellPhoneDB v5: inferring cell-cell communication from single-cell multiomics data. arXiv; 2023. doi:10.48550/arXiv.2311.04567

