## Supplementary figures and images for "Evaluation of Generative AI Models for Processing Single-Cell RNA-Sequencing Data in Human Pancreatic Tissue"

### S1 Appendix

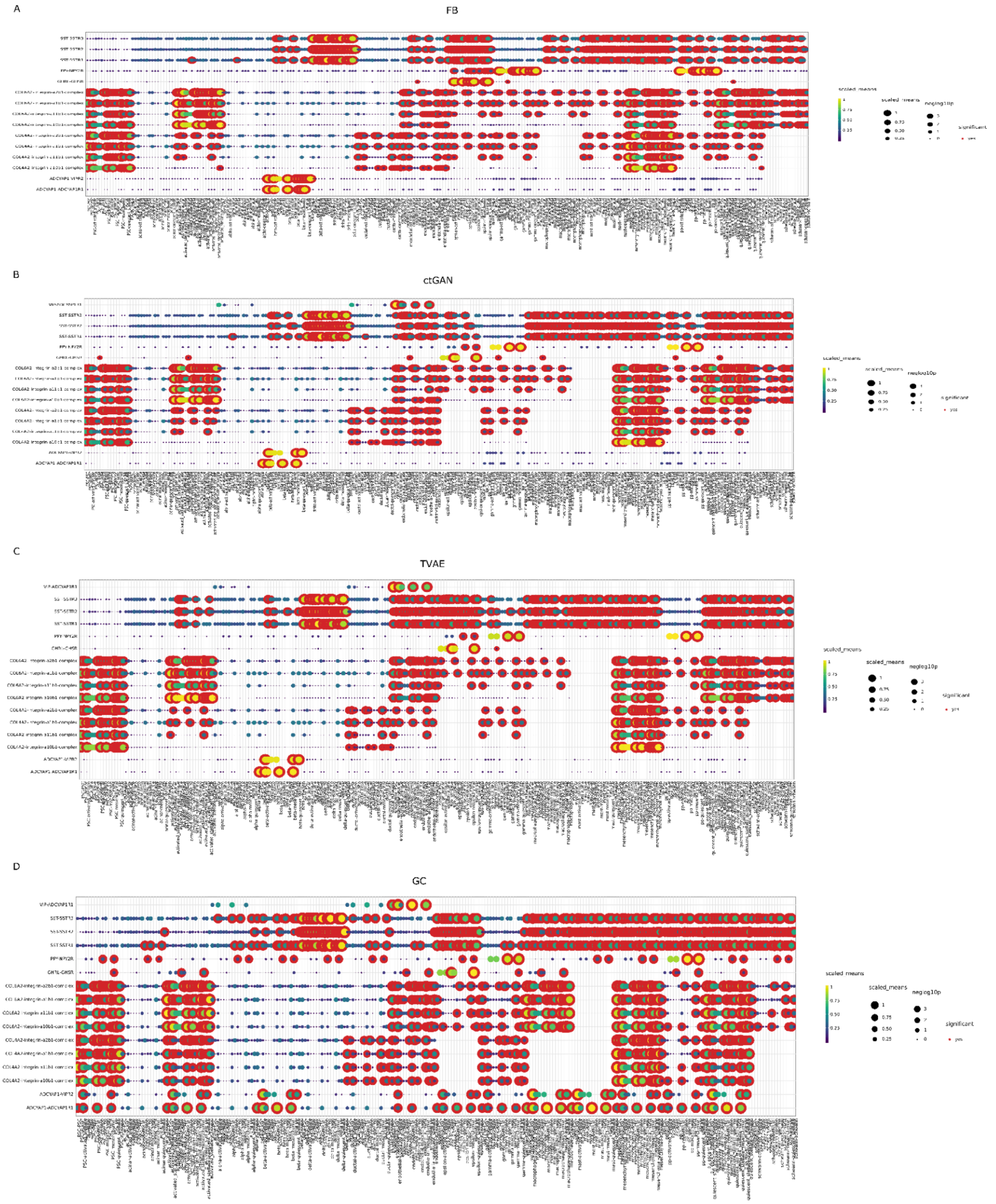


S1 Ligand-receptor pairs of cell-cell interactions (A) FB, (B) ctGAN, (C) TVAE, (D) GC.
